# eSIG-Net: Accurate prediction of single-mutation induced perturbations on protein interactions using a language model

**DOI:** 10.64898/2026.03.27.714913

**Authors:** Xingxin Pan, Aditya Shrawat, Sidharth Raghavan, Chuanpeng Dong, Yuntao Yang, Zhao Li, W. Jim Zheng, S. Gail Eckhardt, Erxi Wu, Juan I. Fuxman Bass, Daniel F. Jarosz, Sidi Chen, Daniel J. McGrail, Gloria M. Sheynkman, Jason H. Huang, Nidhi Sahni, S. Stephen Yi

## Abstract

Most proteins exert their functions in complex with other interactors. Single mutations can exhibit a profound impact on perturbing protein interactions, leading to human disease. However, predicting the effect of single mutations on protein interactions remains a major computational challenge. Deep learning, particularly protein language models or transformers, has become an effective tool in bioinformatics for protein structure prediction. However, the functional divergence of mutations makes it difficult to predict their interaction perturbation profiles. To address this fundamental challenge, we present **eSIG-Net** (edgetic mutation Sequence-based Interaction Grammar Network), a novel sequence-based “Interaction Language Model” for predicting protein interaction alterations caused by single mutations. eSIG-Net combines various protein sequence embeddings, introduces a mutation-encoding module with syntax and evolutionary insights, and employs contrastive learning to evaluate mutation-induced interaction changes. eSIG-Net significantly outperforms current state-of-the-art sequence-based and structure-based prediction methods at predicting mutational impact on protein interactions. We highlight examples where eSIG-Net nominates causal variants with high confidence and elucidates their functional role under relevant biological contexts. Together, eSIG-Net is a first-in-kind “interaction language model” that can accurately predict interaction-specific rewiring by single mutations with only sequence information, and exhibits generalizability across biological contexts.

## Introduction

Substantial improvements in genome and exome sequencing technology in the past 15 years have identified a surfeit of human genetic variation orders of magnitude more extensive than what was previously appreciated. However, how most variants influence the molecular properties and functions of molecules they encode, as well as their impacts on disease initiation and progression remain largely unknown^1^. Among these genetic variants, missense variants are the most common type of protein-coding mutations. Even single missense variants can drastically change protein-protein interactions (PPI)^2, 3^, and therefore rewire protein signaling^4^. Similar to the “activity cliff”^5^ problem in chemistry machine learning, where small structural changes often lead to large or unpredictable changes in activity, single mutations pose an “interaction cliff” grand challenge, causing computational models to mispredict mutation-mediated PPIs.

Machine learning methods have been extensively utilized to characterize protein properties, such as in the prediction of biological functions, protein-protein interactions, and mutational impact on protein stability. Some models utilize amino acid sequences to predict PPIs, such as SDNN^6^, D-SCRIPT^7^, DeepFE^8^, PIPR^9^ and PLM-interact^10^. However, these sequence-based methods do not explicitly model perturbations caused by single amino acid substitutions. Such approaches take the entire protein sequence into account, with the assumption that wild-type and mutant proteins would generate similar embedding vectors in the “protein space”. Therefore, for both wild-type and mutant, these methods may predict similar patterns of protein properties, thus leading to inaccuracy in differential function predictions.

Other methods have shown the potential of using protein structures in predicting impact of missense variants on PPIs, such as MutaBind2^11^, BeAtMuSiC^12^, GeoPPI^13^, TopNetTree^14^ and PIONEER^15^, but all of them require structures of protein complexes as input. This, in turn, limits the application scope of these tools due to scarcity of protein complex structures. Another drawback of structure-based methods is that the structure data often contains ambiguity due to the complexity and variability of experimental conditions.

Applying protein language models is a potential solution to these limitations and has been implemented in methods such as ESM1b^16^, ESM-2^17^, ProtT5^18^, ESM3^19^, D-SCRIPT^7^, and AlphaMissense^20^. However, these methods also face at least two significant challenges. First, they do not explicitly learn the sequence distinctions between mutant proteins and their corresponding wild-type counterparts. Second, they fail to capture the inherent complexity of protein-protein interactions, which are critical for PPI-related tasks.

Contrastive learning is a type of machine learning approach that aims to learn useful representations of data by bringing similar examples closer in the learned representation space while pushing dissimilar examples apart^21, 22^. Compared to canonical representation learning methods, contrastive learning enhances feature representations and leads to more robust models, as it focuses on capturing the nuanced differences between data points. This approach has demonstrated significant success in improving generalization, particularly in tasks such as image recognition and natural language processing, where contextual understanding and subtle distinctions are crucial^23, 24^. More recently, contrastive learning has been effectively applied in proteomics, facilitating tasks like protein structure prediction, protein-protein interaction analysis, and functional annotation by leveraging the discriminative power of contrastive representations^25–27^. This growing body of work highlights the versatility and effectiveness of contrastive learning across diverse biological data types.

Here we introduce a novel interaction language model named **eSIG-Net** (edgetic mutation Sequence-based Interaction Grammar Network). This framework uses the multiple protein embeddings to represent various protein properties^17, 28–30^, leverages a mutation-encoding module to learn the differences in the embeddings of mutation sites, and incorporates a constrained discrepancy module to accurately profile the perturbation of specific PPIs by a given mutation. We benchmark by comparing to current state-of-the-art sequence-based and structure-based PPI prediction methods. Remarkably, eSIG-Net significantly outperforms all these methods at predicting held-out variants for their impact on PPIs. We highlight examples where eSIG-Net nominates causal variants with high confidence and elucidates their functional role under relevant biological contexts. Together, eSIG-Net is a first-in-kind generalizable “interaction language model” that can accurately predict interaction-specific rewiring by genetic mutations or natural variants with only sequence information.

## Results

### The overall framework of eSIG-Net

Conventional protein-protein interaction network prediction involves encoding protein sequences and potential interactors separately, merging these representations, and using a predictor to determine binding states. In this conventional PPI prediction framework, there are typically two modules, namely Encoder 1 and Encoder 2 (**Supplementary Fig. 1**), that process protein sequences individually. Their resulting embeddings are then concatenated and passed through a merged layer. A predictor subsequently uses these merged embeddings to predict interactions. However, determining mutational impact on protein interactions becomes a major challenge due to the minimal differences between mutant and wild-type proteins.

In contrast to conventional PPI prediction methods, eSIG-Net focuses on the discrepancy between wild-type and mutant proteins, as well as their PPI profiles with a specific interaction partner. As shown in **Fig. 1a**, the framework of eSIG-Net consists of two encoder modules: (1) The first is a PPI “protein encoder” module, which is commonly employed in classical PPI prediction tasks. It typically involves separately obtaining the encodings of a protein and its interactor and then merging them to predict PPIs. In the PPI perturbation prediction pipeline, we obtained the merged encodings of both the wild-type with its interactor and the mutant with its interactor. These merged encodings were then fed into a constrained discrepancy module to attempt to discern the differences between them. (2) The second module is a mutant “protein language model” encoder. Extensive empirical evidence has demonstrated that leveraging protein language models could capture the evolutionary information of proteins, thereby facilitating various downstream protein-related tasks. To accentuate the differences between mutant and wild-type proteins, we exclusively utilized the residue-level embeddings of the mutation sites. This was processed through channel-wise learning to obtain a merged mutation site encoding (**Fig. 1a**). Finally, the two merged encodings were integrated and fed into a discriminator for the prediction of PPI perturbations. Compared with the conventional PPI prediction methods, eSIG-Net thus employs an innovative discrepancy strategy to effectively discern the effects of single amino acid changes on proteins and predict ensuing perturbations in their interaction profiles.

**Figure 1.**
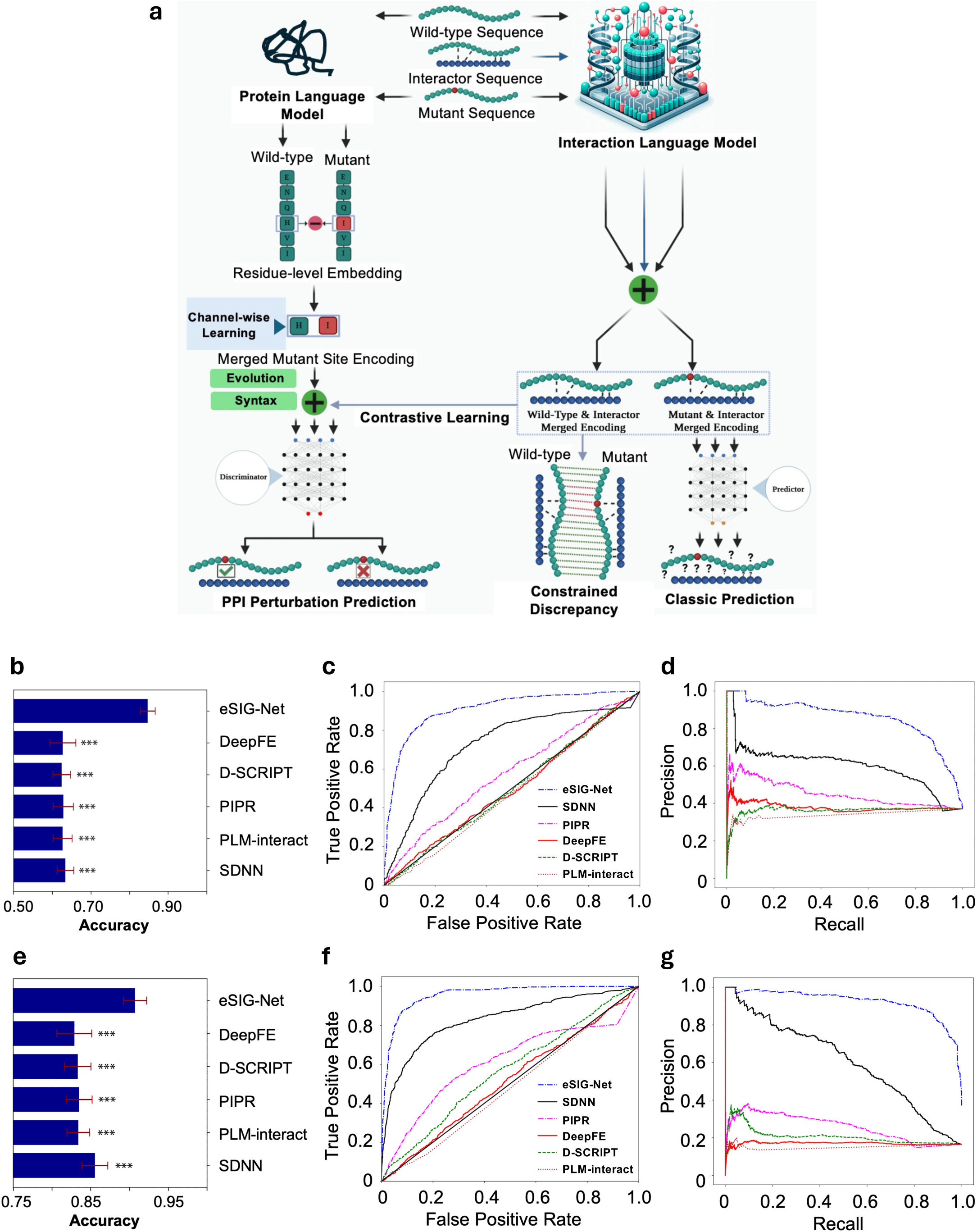
Overview of the eSIG-Net model, and benchmarking with state-of-the-art sequence-based prediction methods. **(a)** The prediction framework of eSIG-Net: wild-type and mutant sequences are processed by a protein language model to obtain residue-level embeddings. These embeddings are then merged through mutation encoding and passed through a channel-wise mutation-site learning module. Concurrently, wild-type/interactor PPI and mutant/interactor PPI pairs are encoded by protein encoder, and their merged encodings are utilized for both constrained discrepancy assessment and traditional PPI prediction. Finally, the combined encodings—mutation-site, wild-type/interactor PPI, and mutant/interactor PPI—are input into a discriminator to predict potential PPI perturbations caused by the mutation. **(b)** Prediction accuracy comparison for the disease mutation PPI dataset, showcasing the performance of eSIG-Net against other PPI prediction models, with statistical significance denoted by asterisks. **(c)** Receiver Operating Characteristic (ROC) curves for the disease mutation dataset, comparing the Area Under Curve (AUC) metrics for eSIG-Net and other models, highlighting eSIG-Net’s superior performance. **(d)** Precision-Recall (PR) curves for the disease mutation PPI dataset, with eSIG-Net outperforming other models in terms of both precision and recall. Line color scheme is the same as (c). **(e)** Prediction accuracy comparison for the gnomAD/ExAC population variant PPI dataset, with eSIG-Net achieving the highest accuracy. **(f)** ROC curves for the population variant PPI dataset, with AUC values for each model, indicating that eSIG-Net maintains a high performance on this dataset as well. **(g)** PR curves for the ExAC population variant PPI dataset, detailing the precision and recall performance of each model, with eSIG-Net providing a competitive precision-recall balance. Line color scheme is the same as (f). *P* values are calculated by pairwise *t*-tests, with Holm-Bonferroni correction. ***, *P* < 0.001.

### Benchmarking PPI perturbation prediction performance

We first benchmarked the performance of eSIG-Net against other methods using two independent datasets: disease mutation PPI dataset (Sahni *et al.)*^2^ and population variant PPI dataset (Fragoza *et al.*)^31^ (see *Methods* for details). Sahni *et al.* dataset is the most comprehensive study (to date) of disease mutations and their effects on PPI profiles. This dataset contains 527 disease mutations in 220 genes, associated with 606 perturbed PPIs and 1,027 non-perturbed PPIs. On the other hand, Fragoza *et al.* dataset carries one of the largest compilations of variants observed in the general population (from gnomAD database), serving as a baseline for neutral or non-pathogenic variation. Fragoza *et al.* dataset contains 1,650 population variants in 772 genes, with a total of 663 perturbed PPIs, and 3,357 non-perturbed PPIs. For each of the datasets, we applied a five-fold cross-validation strategy to avoid the influence of random samples on the performance.

Most existing methods are not specifically tailored to predict PPI perturbations caused by missense mutations, therefore we compared our eSIG-Net model against four state-of-the-art sequence-based PPI prediction methods: SDNN^6^, D-SCRIPT^7^, DeepFE^8^, PIPR^9^, and PLM-interact^10^ (see *Methods* for details). As described above, the eSIG-Net model predicts PPI alterations by mutations, involving triplets composed of the wild-type (WT) protein, the mutant (MT) protein, and the interactor, with the output indicating whether the mutation results in a specific interaction change. In contrast, conventional PPI methods (such as SDNN^6^, D-SCRIPT^7^, DeepFE^8^, PIPR^9^, and PLM-interact^10^) typically take protein-interactor pairs as input, predicting the binding state of the protein-interactor pair. To align inputs for a fair comparison, we split the triplets into WT-interactor and MT-interactor pairs, treating them as two separate samples for conventional PPI tasks. For performance evaluation, we compared the predicted binding state of MT-interactor and WT-interactor pairs, to infer whether the mutation led to a change in interactions.

We first computed the “accuracy” metric, which assesses how correct when predicting perturbed vs unperturbed interactions by a model or method. As shown in **Fig. 1b**, eSIG-Net significantly outperformed all the benchmarking methods, and achieved an accuracy improvement of more than 20% compared to the other methods on the disease mutation PPI data (eSIG-Net accuracy = 0.85; best accuracy by other methods = 0.63). Due to the limited sample size and extremely high similarities between wild-type and mutant proteins, even state-of-the-art PPI prediction methods could not achieve an accuracy above 70%. The results indicate that focusing on the distinct learning of mutation sites enables the recognition of potential missense mutation patterns and their dependencies on interactors. In addition, eSIG-Net outperformed the other existing methods in ROC curve analysis. eSIG-Net achieved an AUC of 0.91, while SDNN had an AUC of 0.73, D-SCRIPT had an AUC of 0.50, DeepFE had an AUC of 0.50, PIPR had an AUC of 0.58, and PLM-interact had an AUC of 0.48 (**Fig. 1c**). Similarly, eSIG-Net also exhibited a better performance in precision-recall curve analysis. eSIG-Net achieved an average precision of 0.86, whereas SDNN had an average precision of 0.60, D-SCRIPT had an average precision of 0.36, DeepFE had an average precision of 0.38, PIPR had an average precision of 0.45, and PLM-interact had an average precision of 0.36 (**Fig. 1d**). These results demonstrate eSIG-Net’s superiority in learning the distinctions between wild-type and mutant proteins, compared to traditional PPI prediction methods that heavily rely on sequence learning (**Supplementary Table 1**).

Next, we assessed the robustness of eSIG-Net in scenarios with severely imbalanced distributions of positive and negative samples (*i.e.*, perturbing PPI or not). This is particularly relevant because in real-world situations, only a very small fraction of missense mutations lead to PPI perturbations or even disease^32^. Over 50% of disease-causing mutations perturb PPIs, while common population variants rarely cause PPI perturbations. We therefore ran our eSIG-Net model on the gnomAD/ExAC population variants PPI dataset. As depicted in **Fig. 1e**, the performance of eSIG-Net remained superior to state-of-the-art PPI prediction methods, achieving an accuracy of 0.90. In contrast, the best accuracy achieved by other methods was 0.86. These results demonstrate that even in scenarios with highly imbalanced positive and negative samples, our method continues to perform exceptionally well. In addition, we performed ROC curve analysis and found that eSIG-Net achieved an AUC of 0.96, while SDNN had an AUC of 0.83, D-SCRIPT had an AUC of 0.56, DeepFE had an AUC of 0.51, PIPR had an AUC of 0.61, and PLM-interact had an AUC of 0.48 (**Fig. 1f**). Similarly, in precision-recall curve analysis, eSIG-Net achieved an average precision of 0.92, whereas SDNN had an average precision of 0.60, D-SCRIPT had an average precision of 0.21, DeepFE had an average precision of 0.17, PIPR had an average precision of 0.26 and PLM-interact had an average precision of 0.16 (**Fig. 1g**). Together, these results indicate that eSIG-Net outperforms other approaches as judged by multiple benchmarks. It’s worth noting that most datasets with variants of uncertain significance (VUS) may be imbalanced. The advantages of eSIG-Net are especially strong for imbalanced datasets, where the limitations of existing methods are even more pronounced (**Supplementary Table 2**).

### Improved interpretability of eSIG-Net model

Existing PPI prediction frameworks can only predict protein-protein interactions individually, and due to the inherent nature of machine learning, they often group similar samples into the same category, making it challenging to distinguish similar samples with different labels. In contrast, eSIG-Net directly learns the differences between samples in a latent subspace.

For a more intuitive comparison, we employed t-distributed Stochastic Neighbor Embedding (t-SNE) to visualize the protein representations generated by the last layer of eSIG-Net and other benchmarking methods. By contrast to the eSIG-Net framework for PPI-perturbation prediction, the other methods required separate inputs of "Mutant/interactor PPI" and "Wild-type/interactor PPI" to obtain their final layer outputs encoding for each triplet. Then we subtracted the "Mutant/interactor PPI" and "Wild-type/interactor PPI" interaction encodings and compared them with the encoding output by eSIG-Net. In the latent space, the encodings generated by eSIG-Net method exhibited much better separation between interaction-perturbing and non-perturbing samples compared to the PPI prediction benchmarking methods (**Supplementary Fig. 1b-g**), using both separation ratio (**Supplementary Fig. 1h**) and silhouette score metrics (**Supplementary Fig. 1i**).

### Evaluating the contributions of individual modules on eSIG-Net prediction

To validate the effectiveness of two main modules in the eSIG-Net framework, we designed and executed an ablation study. In downstream tasks involving protein sequences, performing 1D-pooling on embeddings extracted from pre-trained protein language models is common to maintain input dimensions and reduce computational complexity. This approach captures contextual information from the entire protein sequence, leading to improved predictions. However, for our task of predicting PPI perturbation caused by missense mutations, the protein sequences before and after the mutations are highly similar. Extracting raw embeddings from the protein language model for both sequences could introduce significant redundancy. In the eSIG-Net framework, we exclusively utilized feature vectors from the mutation sites.

As a baseline control, we performed traditional ESM^17^ pooling (esm2_t33_650M_UR50D) embeddings (“Standard Models”, **Fig. 2a**), which yielded a limited improvement in accuracy (0.69; **Fig. 2a**) compared to other benchmarking methods (**Fig. 1b**) on the disease mutation PPI dataset. Moreover, the accuracy on the imbalanced population variant PPI dataset (0.72; **Fig. 2b**) was even lower than the worst-performing benchmarking method (**Fig. 1e**). However, incorporating our mutation site encoding module led to accuracy improvement on both datasets, reaching 0.75 and 0.78, respectively (**Fig. 2a-b**). Finally, with the introduction of our constrained discrepancy learning module, the model’s performance observed further enhancement (accuracy on disease mutation dataset = 0.85, accuracy on population variant dataset = 0.90) (**Fig. 2a-b**).

**Figure 2.**
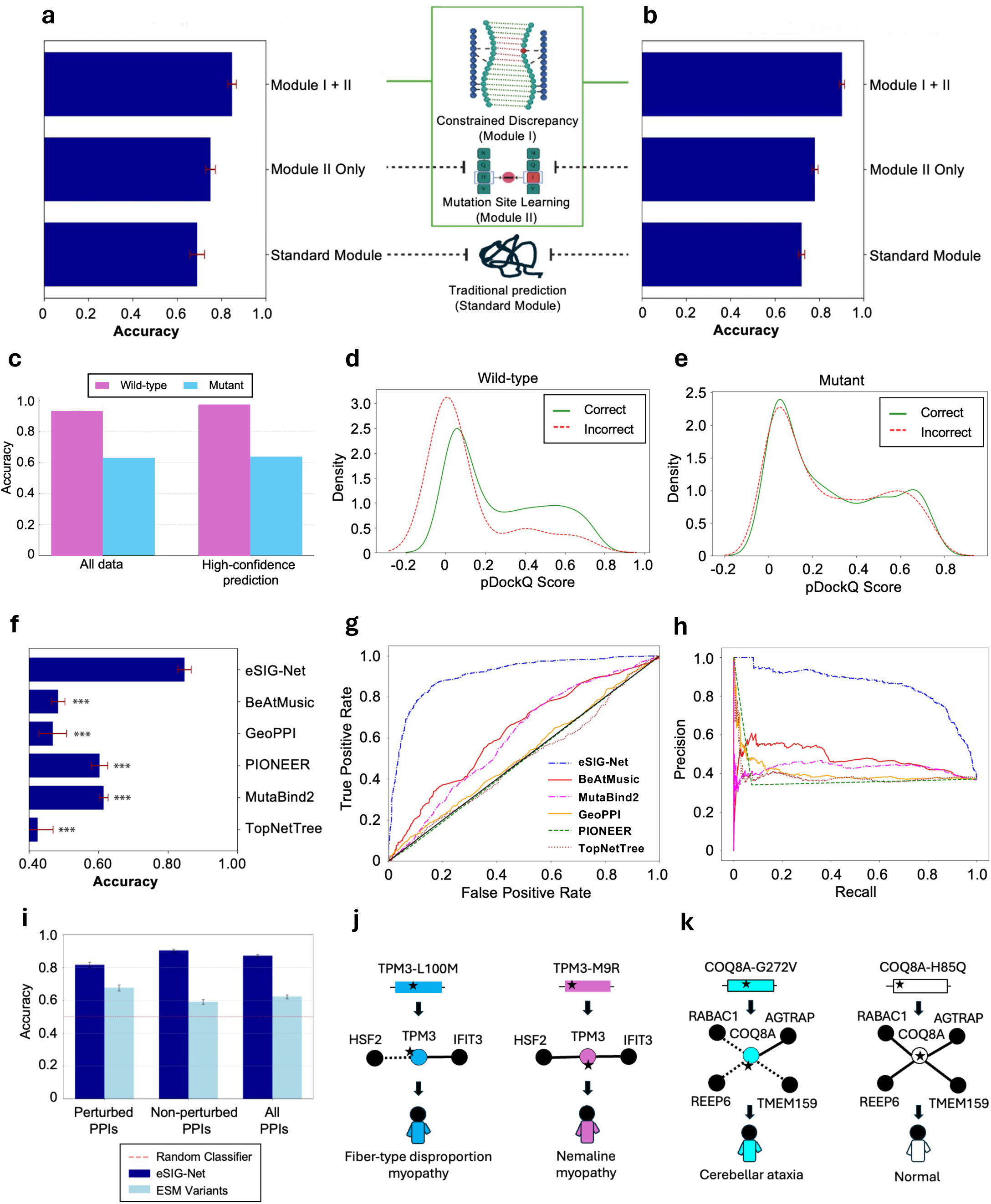
Benchmarking eSIG-Net with mutation-centric, structure-based prediction tools, and application across diverse biological contexts. **(a-b)** Our ablation study showcasing mean accuracy on the disease mutation PPI dataset **(a)** and the population variant PPI dataset **(b)**. Error bars denote the standard errors of the mean. **(c)** Bar chart summarizing the mean accuracy of FoldDock for predicting wild-type vs. mutant protein interactions using all data (left) or high-confidence predictions (pDockQ>0.5; right). **(d)** Density distribution plot of pDockQ scores with correct (green) and incorrect (red) cases, for wild-type protein-mediated interactions. **(e)** Density distribution plot of pDockQ scores with correct (green) and incorrect (red) cases, for mutant protein-mediated interactions. **(f-h)** Prediction evaluation using mutation-centric, structure-based methods: Wild-type and interactor sequences are fed into the AlphaFold-Multimer model, which includes MSA searching and prediction of the complex structure. Then, the mutation information and predicted structure are inputted into the mutation-centric structure-based prediction model to determine the change in binding affinity caused by a mutation. Disease mutation PPI dataset is used here. **(f)** Bar chart summarizing the mean accuracy of different structural algorithms, with the length of each bar representing the mean accuracy and the error bars denoting the standard errors of the mean. **(g)** ROC curves displaying the comparative AUC values for various structural algorithms. **(h)** Precision-Recall curves for structural algorithms. **(i)** Bar chart showing the mean accuracy of ESM-variants (cyan; a mutation-centric disease-causing prediction tool), as compared to eSIG-Net (blue). Bar length represents the mean accuracy and error bars denote the standard errors of the mean. Dashed line indicates a random classifier. **(j)** eSIG-Net-predicted interaction profiles of two disease mutations in the pleiotropic gene TPM3. **(k)** eSIG-Net-predicted interaction profiles of a disease mutation and a population variant in the gene COQ8A (ADCK3). *P* values are calculated by pairwise *t*-tests, with Holm-Bonferroni correction. ***, *P* < 0.001.

### AlphaFold based PPI prediction tool evaluation

FoldDock^33^ applies the AlphaFold^34^ pipeline to simultaneously fold and dock protein-protein pairs, achieving state-of-the-art performance on PPI prediction tasks. Additionally, the model distinguishes high-confidence PPI predictions using a predicted DockQ score (pDockQ). Although FoldDock is a PPI prediction method based on MSA information, different from our sequence-only based method, we explored the potential and constraints of applying the FoldDock prediction tool to simulate interactions between mutant (or wild-type) protein and their interactors.

Using the germline disease mutation dataset (**Supplementary Table 3**), FoldDock’s predictions for wild-type/interactor PPI PPIs were highly accurate, with an accuracy of 0.922, and an even higher accuracy of 0.971 for data with high-confidence scores (pDockQ>0.5), aligning with the conclusions of the original paper^33^. However, the prediction accuracy for mutant/interactor PPI pairs dropped to 0.625, and only 0.632 even with high-confidence predictions (**Fig. 2c**). Moreover, we observed that FoldDock’s predictions for PPIs involving a mutant and its interactor, as well as the corresponding wild-type and its interactor, are largely consistent. Specifically, in 94.4% of the samples, the prediction for a specific PPI involving a mutant protein or its wild-type counterpart remained the same (**Fig. 2d** and **2e**). Furthermore, within the subset of samples where predictions were inconsistent (5.6%), only 17.6% were accurately predicted as disruptive of protein interactions (**Fig. 2d** and **2e**). Therefore, the FoldDock method appears incapable of discerning the subtle differences in proteins with versus without missense mutations, nor can it accurately predict the perturbations in PPI states induced by such mutations.

### Benchmarking with mutation-centric, structure-based PPI prediction methods

Mutation-centric, structure-based PPI prediction methods can be categorized into two main groups: 1) Methods based on physical quantities as input features, such as MutaBind2^11^ and BeAtMuSiC^12^, which introduced several features including interactions of proteins with the solvent, evolutionary conservation, and the thermodynamic stability of complexes for predicting binding changes upon mutations. The reliance on features derived from known rules in protein structures often limits their prediction’s generalization across various samples due to the specificity of these features. 2) Methods based on geometric representation or topological information, such as GeoPPI^13^, TopNetTree^14^ and PIONEER^15^, which circumvent the limitations of manually engineered input features. These approaches significantly accelerate the inference process by utilizing direct representations of protein structures. However, it still necessitates the complete structure of protein complexes as input and requires accurate, experimentally determined binding affinities as labels for training the machine learning models. Despite these different requirements, we further compared the performance of our eSIG-Net model with these two categories of structure-based methods.

Since structure-based prediction tools require the input of protein complex structures, we first subjected the protein sequences to the AlphaFold-Multimer model^35^ to predict their structures, which then served as input for five structure-based prediction methods (MutaBind2, BeAtMuSiC, GeoPPI, TopNetTree and PIONEER). For comparative purposes, mutations classified by these methods as deleterious were considered to be interaction-perturbing (see *Methods* for details) (**Supplementary Table 4**). All of the five benchmarking prediction tools only had an around or below 60% accuracy rate (**Fig. 2f** and **Supplementary Fig. 2a**), much lower than eSIG-Net. In ROC curve analysis, eSIG-Net achieved an AUC of 0.91, while MutaBind2 had an AUC of 0.61, BeAtMuSiC had an AUC of 0.63, GeoPPI had an AUC of 0.52, TopNetTree had an AUC of 0.49, and PIONEER had an AUC of 0.49 (**Fig. 2g**). Similarly, eSIG-Net also exhibited a better performance in precision-recall curve analysis. eSIG-Net achieved an average precision of 0.86, whereas MutaBind2 had an average precision of 0.43, BeAtMuSiC had an average precision of 0.48, GeoPPI had an average precision of 0.40, TopNetTree had an average precision of 0.39, and PIONEER had an average precision of 0.37 (**Fig. 2h**). Together, we found that eSIG-Net significantly outperformed all the mutation-centric structure-based PPI prediction benchmarking tools. A potential reason for this could be that these methods require highly accurate experimental protein complex structures as input, while predicted structures may introduce some level of noise. This can be detrimental when predicting the effects of mutations. Furthermore, despite leveraging advanced machine learning technologies, these tools do not sufficiently differentiate between highly similar samples. In particular, they do not address the intricate challenge posed by missense mutations, which can introduce subtle but critical differences.

### Comparison with ESM scoring methods

The ESM1b method^16^ calculates LLR (Log-Likelihood Ratio) scores for all potential missense mutations within a protein, achieved in a single neural network pass. Using the wild-type (WT) amino acid sequence as input, ESM1b produces the log-likelihood values for each of the 20 standard amino acids, including the WT amino acid, at every position along the protein sequence. The LLR score for each mutation is then determined as the difference between the log-likelihood of the missense amino acid and that of the WT amino acid at the corresponding position. However, this direct calculation of LLR scores from ESM embeddings only provides a reference probability for missense mutations at each position of the WT sequence. It cannot incorporate information from interactors, making it unsuitable for binary classification of PPI perturbations. As a proxy, we normalized the ESM1b scores and used them as confidence scores for PPI perturbations, which were then compared with the logits scores output by the eSIG-Net model (**Supplementary Table 5**). As shown in **Fig. 2i**, eSIG-Net model’s confidence scores exhibited a stronger correlation with the ground truth labels (*i.e.*, PPI perturbation or not) compared to the ESM1b method. Although our mutation site encoding module also relies on embeddings generated by the protein large language model, our framework is trainable and specifically focuses on learning amino acid embeddings for mutation sites. Moreover, with the help of the constrained discrepancy learning and the discriminator, our model can distinguish between the same missense mutations when confronted with different interactors.

### eSIG-Net accurately predicts interaction perturbations, revealing novel mechanistic insights in relevant biological contexts

Although millions of coding variants have been identified in the human genome, the vast majority of them remain classified as variants of unknown significance (VUS). To address this challenge, eSIG-Net offers a generalizable framework that can be applied across diverse biological contexts and adapted to predict interaction-specific variant effects directly from protein sequence. The eSIG-Net model trained on PPI datasets derived from both disease-associated and population-based variants was employed to predict the impact of new (previously ‘unseen’) mutations on PPI perturbation. The training data were partitioned into five distinct sequence-based subsets, each serving as a fold in cross-validation. To ensure a stringent assessment, test set proteins were completely different from those in the training sets. This model demonstrates consistently robust performance in prediction across different sequence groups.

It is important to note that, current state-of-the-art sequence-based and structure-based methods including AlphaFold^34^-derived FoldDock^33^ are limited, and failed to predict interaction alterations by select disease mutations. Remarkably, the eSIG-Net model was able to accurately predict specific PPI perturbations by these mutations. A good example is pleiotropism, where different mutations in the same gene cause different diseases. In the pleiotropic gene TPM3, two mutations L100M and M9R cause fiber-type disproportion myopathy^36^ and nemaline myopathy^37^, respectively (**Fig. 2j**). Intriguingly, eSIG-Net predicted the mutation L100M to selectively perturb (*i.e.*, edgetic) the interaction with HSF2, which was known to expressed in muscle and involved in myotube regeneration^38^. In contrast, eSIG-Net predicted the mutation M9R to retain the interaction with HSF2 (**Fig. 2j**). Taken together, eSIG-Net provided possible mechanistic insights into pleiotropic phenotypic outcomes through accurate prediction of distinct interaction profiles.

As another example, COQ8A (ADCK3) encodes a mitochondrial protein that functions in an electron-transferring membrane protein complex, and the mutation G272V in COQ8A leads to cerebellar ataxia^39^. eSIG-Net correctly predicted that the COQ8A-G272V mutation impaired the interactions with RABAC1 (intracellular trafficking protein involved in neuropathy), REEP6 (transport of receptors from ER to cell membrane in neural development), and TMEM159 (LDAF1) (formation of lipid droplets in the ER) (**Fig. 2k**). On the other hand, eSIG-Net predicted the population variant COQ8A-H85Q to preserve all these interactions (**Fig. 2k**). In summary, this likely represents a false-negative case by current state-of-the-art methods, but instead is accurately predicted as disruptive of PPIs by the eSIG-Net method.

Further analysis of public genomics data revealed that the majority of eSIG-Net-predicted PPI pairs mediated by mutations (**Supplementary Table 6**) were associated with clinical outcomes. Specifically, 46.8% of these pairs were linked to prognosis in more than five cancer types across The Cancer Genome Atlas (TCGA) and the Moffitt Cancer Center (MMRF) cohorts (**Supplementary Fig. 2b**). We also observed that several predicted PPIs also significantly correlated with responses to cancer immunotherapy (**Supplementary Fig. 2c**).

Together, these results underscore the specific biological contexts captured by eSIG-Net regarding the functional significance of genetic variants in perturbing specific protein interactions, facilitating our understanding of the large numbers of variants of unknown significance. The insights that a given variant impacts specific interactions within signaling pathways provide a critical new perspective to understand pathological mechanisms that would otherwise be missed. This further highlights eSIG-Net’s potentially substantial impact as a vehicle for mechanistic discovery.

## Discussion

Although protein language models are commonly used to extract protein-level embeddings for predictive tasks^17^, they do not sufficiently capture the functional changes (such as interaction perturbations) induced by single mutations in a protein sequence. Similar to the “activity cliff” problem in the chemistry field, this “interaction cliff” concept by single protein mutations poses a major computational challenge. In the pursuit to discern the intricate perturbations of PPI states due to missense mutations, we present eSIG-Net, a state-of-the-art interaction language model and deep learning framework tailored to understand and predict such perturbations. In this method, we introduce a novel discrepancy module to amplify the differences between the PPI-pair encoded representations before and after the mutation. Compared to conventional PPI predictors, we design a discriminator to assess the perturbations in PPI states resulting from the mutation, thereby enhancing the model’s robustness. Finally, we leverage the powerful protein language model ESM-2^17^ to obtain embeddings of the protein sequences before and after the mutation, which serve as additional inputs. The additional inputs are then fed into a uniquely designed mutation site channel-wise encoding module.

One of the key elements of eSIG-Net is to encode wild-type/interactor PPI and mutant/interactor PPI protein pairs separately. Simply differentiating the mutant and wild-type encodings alone is not sufficient, as many missense mutations are edgetic^2^, resulting in a perturbation depending on specific interactors. Besides the separate paired encodings, a mutation site encoding module is introduced to embed mutation-specific information.

To the best of our knowledge, there is currently no method that applies sequence based deep learning technologies to the analysis of PPI perturbation caused by single mutations. Our “Interaction Language Model” eSIG-Net surpasses the existing advanced PPI methods on diverse datasets, indicating its high accuracy and robustness. Notably, our framework offers exceptional versatility, accommodating diverse input features and integrates seamlessly with advanced neural networks, enabling flexible incorporation into emerging technological advancements in deep learning. More importantly, our proposed method fundamentally aims to address the challenging machine learning problem posed by the biological question of PPI perturbation caused by single amino acid changes. This problem involves distinguishing between samples with potentially vastly different labels but subtle differences in protein sequence. This innovative framework brings to the fore a novel mechanism to handle the challenges posed by the inherent properties of missense mutations, which can drastically modify protein-protein interactions even with a single amino acid in the primary sequence.

Traditional approaches to computationally model the changes in binding/interaction states due to mutations typically involve structural modeling techniques^40^ or empirical energy-based methods^41^. But these methods demand substantial computational resources^42^ and/or often struggle with inadequate conformational sampling, particularly for mutations in flexible regions^13^. The advancement of machine learning has presented an opportunity to model the intrinsic relationship between a mutation and its functional effect on protein interaction or binding affinity directly. However, these models often require experimentally determined protein structures, which restricts their broad applicability due to the challenges to obtain precise structural data of protein complexes^43^. This problem is especially acute for intrinsically disordered sequences that adopt distinct conformations when interacting with different protein partners, and are enriched in regulatory factors often mutated in human disease.

Currently, large-scale studies of mutational impact on protein activities are extremely challenging, which are primarily measured by high-throughput wet-lab experimental platforms, such as functional variomics^3^ and deep mutational scanning^44^. Although these methods have made enormous strides in characterizing large numbers of protein variants, they remain time-consuming and labor-intensive. The eSIG-Net method is designed to exactly tackle this problem, and can serve as an accurate and alternative functional characterization of variants at large scale through deep *in silico* mutagenesis. This effort will greatly facilitate the annotation and analysis of many protein variants of unknown significance thus far, potentially contributing to discovery of novel disease-relevant biomarkers and therapeutic strategies.

Similar to other state-of-the-art methods, eSIG-Net also possess potential limitations. First, methods that utilizing Multiple Sequence Alignment (MSA) to extract information typically yield embeddings that capture valuable mutation and evolutionary information^20^. For example, certain general protein structure prediction tasks heavily rely on MSA information^34, 45^. However, the process of extracting MSA data is slow, and demands substantial computational resources. In the eSIG-Net framework, we use sequence-based biostatistical embeddings and protein language model embeddings for both the input to the PPI prediction module and the mutation site encoding module to expedite embedding extraction. Nevertheless, this approach sacrifices some co-evolutionary information under specific biological contexts. While the current version of eSIG-Net primarily predicts mutational effects on the energetic or biophysical favorability of interactions between a pair of proteins, there’s room for future development considering that many disease-causing mutations lead to disease in a tissue-specific manner. Optimization of the eSIG-Net model by incorporating gene expression information as an input would facilitate the prediction of actual interactions within a given ‘disease-relevant’ tissue.

Nor does a change of PPI forcedly reveal causation of disease, let alone the druggable target identification. These goals will require further model development. Nevertheless, we believe that eSIG-Net has the potential to revolutionize our comprehension of the critical effects caused by missense mutations in molecular networks and to catalyze substantial advancements in therapeutic interventions for genetic disorders.

## Methods

### eSIG-Net model

We first assembled multiple effective feature generation methods for PPI prediction to enhance the representation of proteins. Then, we optimized the PPI prediction framework by introducing a constrained discrepancy learning module. This module was specifically designed to differentiate the merged encoding of "mutant/interactor PPI" and "wild-type/interactor PPI" pairs, as their differences are subtle yet crucial for capturing the effects of missense mutations on PPI states. Additionally, we harnessed the capabilities of protein language models by incorporating mutation site encoding into the embeddings obtained from these models. We also employed a discriminator to predict the impact of missense mutations on PPI states.

It’s essential to note that eSIG-Net primarily focused on learning the differences between wild-type and mutant protein embeddings, in conjunction with interactors, to predict the occurrence of PPI perturbations. Rather than directly predicting PPI states, our approach took a distinct strategy. In comparison to traditional PPI prediction methods, eSIG-Net incorporated several innovative designs to effectively distinguish between similar samples (*i.e.*, protein sequences that differ by a single mutation), and predict perturbations in their PPI states.

### Datasets

Two mutation-mediated PPI datasets were used in this study: Sahni *et al.* disease mutation PPI dataset^2^, and Fragoza *et al.* population variant PPI dataset^31^. Sahni *et al.* dataset consists of 1,633 samples, with each sample composed of three proteins (“triplet”: the wild-type protein, the mutant protein, the interactor) and the binding states of the wild-type/interactor PPI (WT-interactor) and mutant/interactor PPI (MT-interactor) pairs (0/1). This dataset contains 527 disease mutations in 220 genes, associated with 606 perturbed PPIs and 1,027 non-perturbed PPIs. On the other hand, Fragoza *et al.* dataset carries one of the largest compilations of variants observed in the general population (from gnomAD database), serving as a baseline for neutral or non-pathogenic variation. Fragoza *et al.* dataset contains 1,650 population variants in 772 genes, with a total of 663 perturbed PPIs, and 3,357 non-perturbed PPIs. We excluded synonymous mutations from this analysis. We further defined a positive sample as one where a PPI state change (*i.e.*, perturbation) occurred, while a negative sample represents cases where no PPI perturbation took place. Note that the population variants dataset was used to validate the robustness of our model and other methods. This dataset contains only approximately 16% positive samples, making it an imbalanced dataset.

To obtain the gene sequences corresponding to the genotypes, we retrieved them from the hORFeome^46^ V9.1 Library (http://horfdb.dfci.harvard.edu/) and converted the nucleotide sequence into amino acid sequences.

### Model input features

In the experiments, the protein sequences were transformed into fixed-length feature vectors before being input into the neural network due to variations in sequence length. The feature fusion strategy was employed to convert protein sequences into feature vectors using three methods: Amino Acid Composition (AAC), Conjoint Triad (CT), and Auto Covariance (AC).

- The AAC method^28^ normalizes the frequency of occurrence of each amino acid in the protein sequence. It counts the frequency of the twenty amino acids, resulting in a 20-dimensional feature vector for each protein sequence.
- The CT method^29^ divides the twenty amino acids into seven different clusters based on the volume of amino acid side chains and dipoles. Each cluster groups together amino acids with similar characteristics. This results in a 343-dimensional feature vector that represents the normalized triples (7*7*7) of amino acids.
- The AC method was used to capture interactions between amino acids that are separated by a specific number of residues within a protein sequence. The process starts by converting the amino acid residues into numerical values that represent their physicochemical properties, such as hydrophobicity, polarity, or molecular weight. After encoding the sequence numerically, the AC calculation is carried out, which involves computing the covariance between the properties of amino acids separated by a fixed distance. The resulting 210-dimensional AC feature vector is normalized to zero mean and unit standard deviation (SD). The experimental setup follows ^6, 30^ for obtaining the AC feature vector.

In summary, these three feature vectors were concatenated to form a 573-dimensional feature vector, which was used for Discrepancy learning and prediction. Additionally, as shown in **Supplementary Fig. 1**, our model utilized ESM-2^17^ to obtain additional embeddings for the mutation site encoding module. Specifically, the “esm2_t33_650M_UR50D” version of the model^17^ was used to obtain residue-level embeddings. We extracted the embedding at the mutation site to obtain a 1,280-dimensional ESM feature vector.

### Constrained discrepancy learning

To achieve the objective of learning a precise amount of discrepancy across mutated proteins with different binding state changes, we defined the distance between the merged encodings of missense mutations before and after as *d_i_* and indicate whether the binding state changes as *c_i_* (*c_i_* = 1 if there is a perturbation, otherwise *c_i_* = 0). We formalized our constrained discrepancy loss in the follow equation (**Equation 1**):

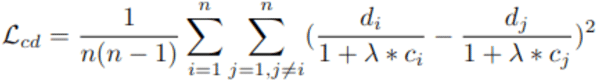

where *n* is the number of the samples in one training batch, and *d_i_* is the embedding-level differences before (*W_i_*, wild-type) and after (*M_i_*, mutant) the mutation, which measured by L2 norm: *d*_*i*_ = | *W*_*i*_ − *M i* |_2_. λ is a hyper-parameter. It is important to note that using **Equation 1** alone is insufficient, as the trivial solution would be to set all the embedding-level distances as zero (*i.e.*, *d_i_* = 0 for all *i*). However, learning the accurate amount of discrepancy while avoiding the trivial solution is achievable by jointly training with the objective of the original PPI prediction change. In this joint training approach, the model was devised to embed "Mutant/interactor PPI" and "Wild-type/interactor PPI" differently to discriminate between them.

### Mutation site encoding

As illustrated in **Supplementary Fig. 1**, we harnessed the capabilities of the protein language model (specifically ESM-2^17^) to acquire residue-level embeddings for both the wild-type (WT) and mutant (MT) proteins. These embeddings capture contextual information from the protein sequence. Subsequently, to reduce redundancy, we isolated the embedding specific to the mutation site and directed it through the mutation site channel-wise self-attention module.

The pipeline of our mutation site encoding module is as follows:

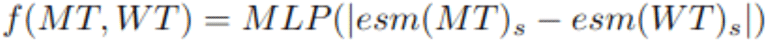

esm(⋅)_*s*_ ∈ *R*^*d*^ represents the vector corresponding to the mutation site extracted from the residue-level embedding (esm(⋅) ∈ *R*^*L*×*d*^), where (L) is the length of the protein and (d) is the feature dimension of ESM. Multi-Layer Perceptron (MLP) was implemented to obtain the final Mutation site encoding.

Subsequently, we concatenated the Mutant/interactor PPI merged, Wild-type/interactor PPI merged, and Mutation site encoding, and fed them into the discriminator to predict whether the missense mutation leads to a perturbation in the interaction state. Our overall learning objective is given as follows:

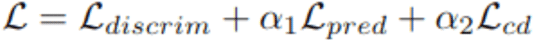

where both the discriminator loss *L_discrim_* and predictor loss *L_pred_* utilize the cross-entropy loss function. The discriminator’s labels are based on whether the interaction is perturbed, with “1” indicating a perturbation and “0” indicating no change. The predictor’s labels represent the interaction itself, where “1” denotes a binding (interaction) and “0” denotes no binding. The hyperparameters α_1_ and α_2_ are used to control the weights of the respective losses. Our model training finally set α1 and α2 to 0.9 and 0.1, respectively.

### Training strategy

The Mutant/Wild-type-encoding module and the Interaction-encoding module shared the same network structure, which consists of seven linear layers forming an MLP. The merge layer was a linear layer with a size of 32. The mutation site encoding module comprised 2 linear layers. Both the predictor and discriminator consisted of two MLP layers serving as classifiers. Batch normalization and a dropout rate of 0.3 were applied after each linear layer in the model. Each of our linear layers (except the output layer) was followed by a ReLU non-linearity, to allow complexity of functions our eSIG-Net model could learn.

For optimization, we employed multi-step learning rate descent with epoch_index = [30] for epoch indices. The learning rate decay factor was set to 0.1. Adam was used as the optimizer for our method, with an initial learning rate of 0.005. The weights α_1_ and α_2_ for the loss function were set to 0.9 and 0.1, respectively. After training 50 epochs, we selected the optimal checkpoint based on the validation accuracy.

### Five-fold cross-validation

We applied a five-fold cross-validation strategy to avoid the influence of random samples on the performance. The eSIG-Net model trained on PPI datasets was employed to predict the impact of new (previously ‘unseen’) mutations on PPI perturbation. The training data were partitioned into five distinct subsets, each serving as a fold in cross-validation. Model selection was performed on the 5th held-out fold. To ensure a stringent assessment, test set proteins were completely different from those in the training sets. This model demonstrates consistently robust performance in prediction across different sequence groups. A use of similarity filter to define the training set and test set was found to make no significant difference, further demonstrating that our model performance was not driven by sequence similarity alone.

### Benchmarking with other sequence-based methods

Due to the absence of existing methods specifically tailored to predict changes in the original protein-protein interaction (PPI) states caused by missense mutations, we sought to compare our framework against four state-of-the-art PPI prediction methods. While both our framework and the benchmarking methods address PPI-related tasks, they differ in their inputs. Our model for predicting changes in PPI states by missense mutations involves triplets composed of the wild-type (WT) protein, the mutant (MT) protein, and the interactor, with the output indicating whether the binding states of WT-interactor and MT-interactor change. In contrast, conventional PPI methods typically take protein-interactor pairs as input, predicting the binding state of the protein-interactor pair. To align inputs for a fair comparison, we split the triplets into WT-interactor and MT-interactor pairs, treating them as two separate samples for conventional PPI tasks.

The logit output of these sequence-based methods is a continuous value between 0 and 1 (with ‘1’ representing the probability of PPI/binding). The binding state of the mutant-interactor was determined by binarizing the logit value. For mutation-perturbed PPIs, the predicted interaction probabilities of the wild-type (WT) PPI are above 0.5, whereas the probabilities of the mutant (MT) PPI fall below 0.5. For non-perturbed PPIs, the predicted interaction probabilities of the WT and MT PPIs are either “both above 0.5” (*i.e.*, both having PPIs), or “both below 0.5” (*i.e.*, both exhibiting no PPIs). For performance evaluation, we compared the predicted binding state of MT-interactor and WT-interactor pairs, to infer whether the mutation led to a change in interactions.

#### SDNN

SDNN^6^ evaluates diverse protein feature extraction combinations and identifies the most effective feature sets. It further enhances performance by employing attention-based networks, achieving results in PPI prediction tasks. In this study, we adopted the optimized feature combinations as reported^6^ and employed their PyTorch codes to train the PPI prediction model.

#### D-SCRIPT

D-SCRIPT^7^ introduces a method for embedding proteins based on their amino-acid sequences, aiming to bring proteins with similar structures closer in the embedded space. It employs a stacked 3-layer Bidirectional LSTM for protein embedding, yielding results in PPI tasks. We followed the same pipeline^7^ and used D-SCRIPT package to train the prediction model.

#### DeepFE

DeepFE^8^ integrates handcrafted features with Word2vec technology, enabling the creation of protein sequence embeddings that capture intricate semantic relationships among amino acids. These embeddings are then harnessed within deep neural networks, proficiently extracting features, reducing dimensionality, and making predictions of protein-protein interactions. In this study, we adopted the original codes^8^ to generate the input embeddings and train the prediction model.

#### PIPR

PIPR^9^ incorporates a Siamese neural network featuring a deep residual recurrent convolutional architecture. This design effectively combines local features and contextual information, enabling the capture of mutual influences within protein sequences. Moreover, PIPR simplifies data pre-processing and demonstrates adaptability across various applications. We utilized the PIPR codes^9^ to train the PPI model.

#### PLM-interact

PLM-interact^10^ leverages joint encoding of protein pairs, enabling the model to directly learn relational patterns between sequences. By fine-tuning pre-trained transformer-based PLMs with contrastive learning objectives, PLM-interact effectively captures both individual sequence features and inter-protein dependencies. In this study, we employed the PLM-interact^10^ implementation to fine-tune the model and predict PPI outcomes.

### Visualization and quantification of model interpretability

Normalized confusion matrix was used for displaying the prediction, and t-distributed Stochastic Neighbor Embedding (t-SNE) plot was used for the visualization of model interpretability. We used two metrics to quantify the degree to which different classes of data points (perturbed vs. unperturbed PPIs) are separated in the visualization: (1) Separation ratio: Separation ratio was computed as the between-centroids distance divided by the root-mean-square within-cluster distance. A higher separation ratio generally indicates that the clusters are more distinct and well-separated in the t-SNE visualization, suggesting that the data points within each cluster are more similar to each other than to points in other clusters. (2) Silhouette score: Silhouette score of each point was computed using the formula: *s* = (*b* – *a*) / max (*a*, *b*), where *a* represents the average distance to all other points within its cluster, and *b* represents the average distance to all points in the nearest cluster. The overall silhouette score is the average of all individual silhouette scores. A high silhouette score indicates well-defined and separated clusters.

### AlphaFold-based PPI prediction

DeepMind’s AlphaFold^34^ offers regular atomic accuracy in protein structure prediction through a multi-sequence alignment (MSA) encoder and build pairwise representations, a 3D rotation-equivariant network to build structure, and an iterative recycling mechanism to optimize structure prediction. This end-to-end structure prediction framework can benefit the analysis of various protein structures, properties, and functions^47^. FoldDock^33^ leverages the AlphaFold^34^ pipeline for protein interaction prediction. To predict mutation-mediated PPI changes, we fed all wild-type/mutant and interactor sequence samples into the FoldDock predictor. As MSA was executed, the quantification of interface contacts was obtained. A count of fewer than one interface contact was interpreted as a non-interaction, or a negative PPI prediction. Given the intensive time requirements for MSA extraction and the substantial GPU resources demanded by the sophisticated AlphaFold2 model, our study’s performance metrics were confined to the disease mutation PPI dataset.

### Benchmarking with mutation-centric, structure-based methods

To compare eSIG-Net with other mutation-centric, structure-based methods, we first subjected the wild-type and interactor sequences to the AlphaFold-Multimer model^35^. This model includes a search for multiple sequence alignments (MSAs) and predicts the structure of the wild-type/interactor PPI complex. With the structures predicted, we then incorporated the mutation information into the prediction model to estimate the changes in binding affinity (ΔΔG) caused by the mutation. Due to these methods classifying a mutation as deleterious if ΔΔG >=1.5 or <= −1.5 kcal/mol, we defined such deleterious mutations as perturbing PPI profiles. To illustrate the AUC curves, we converted (ΔΔG) into a logit score by performing the following operation: score = sigma (| ΔΔG | - 1.5), where (sigma) represents the sigmoid function. Disease mutation PPI dataset was used for benchmarking with all structure-based methods.

### Population-based PPI function validation in the context of cancer immunotherapy

To validate the potential impact of mutations on eSIG-net predicted PPI pairs, we obtained transcriptomic and somatic mutation data from the TCGA and MMRF-COMPASS cohorts via the UCSC Xena platform^48^, encompassing 34 cancer types and over 11,000 patients. For analysis on the response to immunotherapy, patients were stratified into a “both-high” group (expression of both PPI genes ≥ median) versus all others. Statistical significance was assessed using Fisher’s exact test.

### Statistical analysis

To compare the mean accuracies of eSIG-Net against the other models, pairwise independent *t*-tests were conducted. Given the multiple comparisons being made (each algorithm against eSIG-Net), it was necessary to adjust for the increased probability of Type I errors. To this end, we employed the Holm-Bonferroni method for adjusting p-values.

## Data and code availability

All datasets, code and documentation are available at https://github.com/Yilab-texas/eSIG-Net.

## Acknowledgments

The authors would like to acknowledge assistance by Ben Hitz, Zhewei Shen, Khine Lin, and Shengcheng Dong from the DACC of IGVF (Impact of Genomic Variation on Function) Consortium. The work was supported by NIH grants R35GM133658 (to S.Y.), R33CA281919 (to S.Y.), R00CA240689 (to D.J.M.), R35GM137836 (to N.S.), R01HG012366 (to D.F.J.), R35GM128625 (to J.I.F.B.), and U24MH130988 (to W.J.Z.). We would also like to acknowledge the Department of Defense grant W81XWH-22-1-0164 (to W.J.Z.). S.Y. is an Affiliate Member of the NHGRI IGVF Consortium, a Partner Member of the NHGRI GREGoR (Genomics Research to Elucidate the Genetics of Rare diseases) Consortium, and an Associate Member of the NCI Cancer Systems Biology Consortium (CSBC). N.S. was supported by Alfred P. Sloan Scholar Research Fellowship (FG-2018–10723). S.C. was supported by Cancer Research Institute Lloyd Old STAR Award (CRI 4964). S.G.E. would like to acknowledge the Dan L Duncan Comprehensive Cancer Center grant P30CA125123.

## Author contributions

S.Y., X.P. and N.S. conceived the study. X.P., A.S. and S.R. conducted most of the computational modeling and analyses, with input from C.D., Y.Y., Z.L., D.J.M., S.Y., and N.S.. E.W., J.H.H., S.G.E., W.J.Z., J.I.F.B., G.M.S., S.C. and D.F.J. provided intellectual input and constructive feedback. S.Y. and N.S. provided supervision throughout the course of the study. X.P., A.S., N.S. and S.Y. wrote the manuscript with input from S.C., D.J.M., J.I.F.B. and D.F.J.. All authors read and approved the manuscript.

---

**Supplementary Figure 1.**
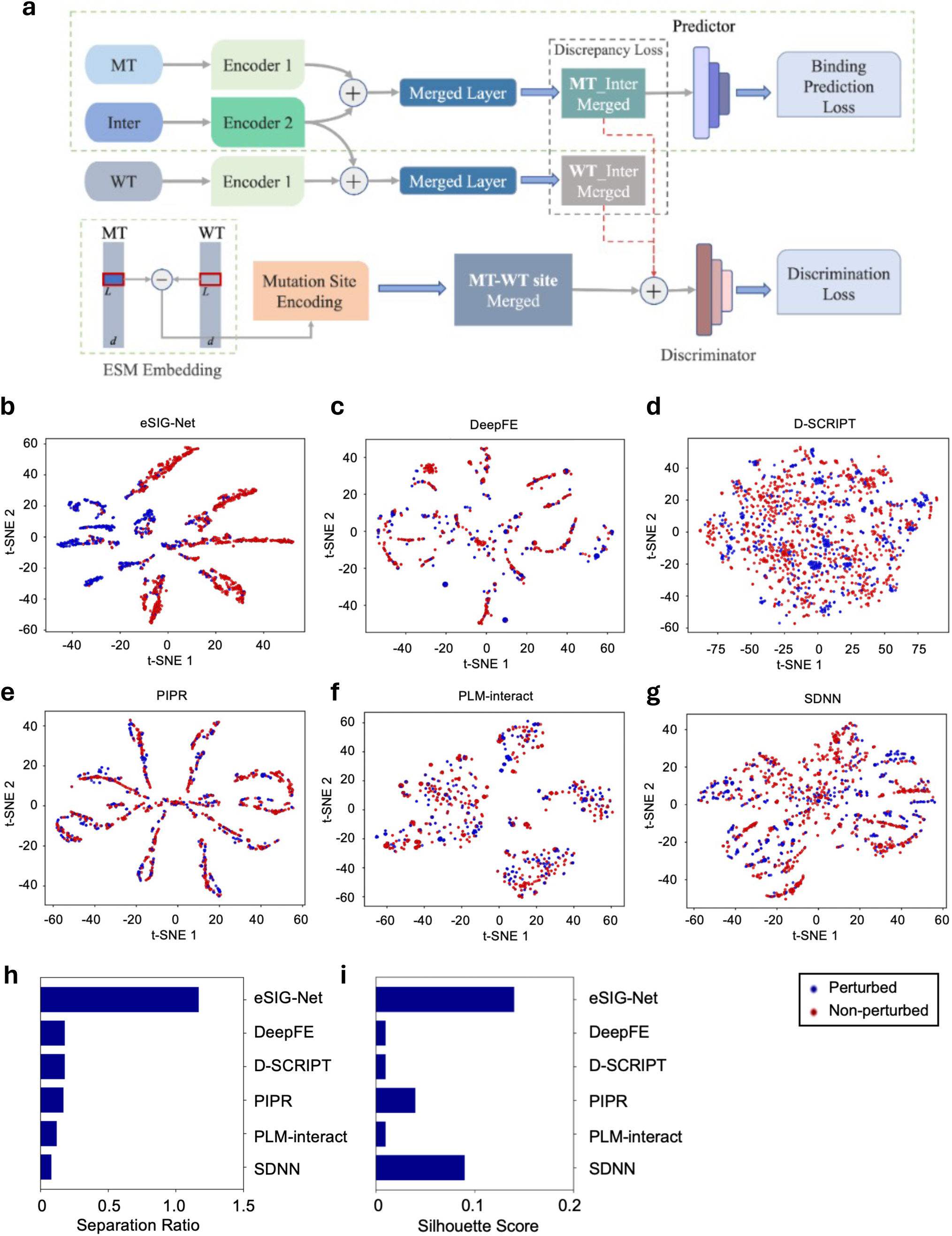
Architecture and interpretability of the eSIG-Net model. **(a)** Two protein encoders are used to encode Mutant (MT)/Wild-type (WT) and Interactor (Inter), with MT and WT encoders sharing weights. The resulting encodings are concatenated and input into the Merged layer, producing MT-Inter merged encoding and WT-Inter merged encoding. Similar to traditional PPI prediction methods, the MT-Inter merged encoding is fed into the predictor for interaction/binding state prediction. Additionally, we calculate the discrepancy loss between MT-Inter merged encoding and WT-Inter merged encoding to constrain the distance between positive and negative samples at the encoding level. We then extract MT and WT embeddings from ESM-2 and subtract the mutation site embeddings to input into the mutation site encoding module to learn differences at the mutation site. Finally, we concatenate MT-Inter merged encoding, WT-Inter merged encoding, and MT-WT site merged encoding and input them into the discriminator for the ultimate PPI perturbation prediction. For details, refer to the *Methods* section. **(b-g)** t-SNE dimensional reduction visualization comparing eSIG-Net (**b**) with other state-of-the-art sequence-based methods, including DeepFE (**c**), D-SCRIPT (**d**), PIPR (**e**), PLM-interact (**f**) and SDNN (**g**). Predicted perturbed and unperturbed PPIs are shown in blue and red color, respectively. Normalized confusion matrix is used for displaying the prediction, and t-distributed Stochastic Neighbor Embedding (t-SNE) plot is used for the visualization of model interpretability. The embedding of the output layer’s previous hidden layer output from all methods is extracted to perform the visualization. For quantification, two metrics (separation ratio and silhouette score) are computed to quantify the degree to which distinct classes of data points are separated in the visualization. **(h)** Barcharts showing the separation ratio metric to compare methods for their prediction performance on separating perturbed vs non-perturbed PPIs, using the disease mutation dataset. **(i)** Barcharts showing the silhouette score metric to compare methods for their prediction performance, using the disease mutation dataset.

**Supplementary Figure 2.**
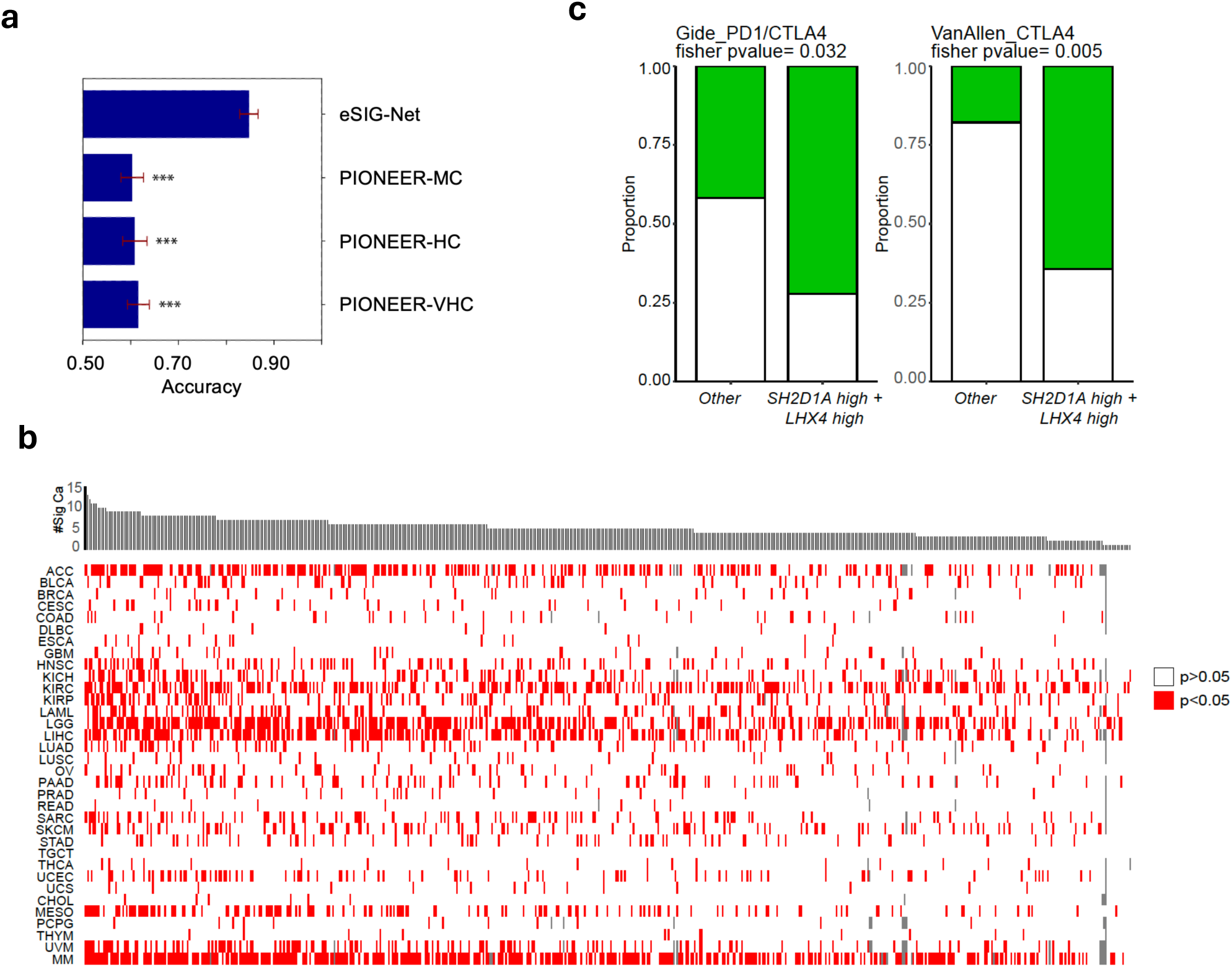
Benchmarking with the state-of-the-art structure-based prediction method PIONEER, and independent validation across different biological contexts. **(a)** Bar chart shows the mean accuracy of the structure-based prediction method PIONEER across different confidence levels (MC: Medium confidence; HC: High confidence; VHC: Very high confidence), using the disease mutation dataset. The length of each bar represents the mean accuracy and the error bars denote the standard errors of the mean. **(b)** PPI-paired genes as predicted by eSIG-Net may jointly influence cancer patient survival in TCGA-MMRF cohorts. Patients are stratified into a "both-high" group (expression of both PPI genes ≥ median) versus all other patients. Cox proportional hazards regression is used to evaluate statistical significance. **(c)** Expression of PPI gene pairs as predicted by eSIG-Net is associated with immunotherapy response in melanoma patients. Gide PD-1/CTLA-4 cohort (*n* = 74); Van Allen CTLA-4 cohort (*n* = 43).

**Supplementary Table 1.** Five-fold validation scores predicted by eSIG-Net, SDNN, D-SCRIPT, DeepFE, PIPR, and PLM-interact using disease mutation PPI dataset.

**Supplementary Table 2.** Five-fold validation scores predicted by eSIG-Net, SDNN, D-SCRIPT, DeepFE, PIPR, and PLM-interact using population variant PPI dataset.

**Supplementary Table 3.** Wild-type and mutant PPI prediction results by FoldDock.

**Supplementary Table 4.** Confidence scores for prediction of mutant-induced PPI perturbations by five structure-based benchmarking methods.

**Supplementary Table 5.** Prediction scores for mutational impact by ESM.

**Supplementary Table 6.** eSIG-net mutation occurrence in TCGA-MMRF cohorts in 34 cancer types

## Reference

1. Ng, P.K. et al. Systematic Functional Annotation of Somatic Mutations in Cancer. Cancer Cell 33, 450–462 e410 (2018).

2. Sahni, N. et al. Widespread macromolecular interaction perturbations in human genetic disorders. Cell 161, 647–660 (2015).

3. Yi, S. et al. Functional variomics and network perturbation: connecting genotype to phenotype in cancer. Nat Rev Genet 18, 395–410 (2017).

4. Li, Y. et al. e-MutPath: computational modeling reveals the functional landscape of genetic mutations rewiring interactome networks. Nucleic Acids Res 49, e2 (2021).

5. van Tilborg, D., Alenicheva, A. & Grisoni, F. Exposing the Limitations of Molecular Machine Learning with Activity Cliffs. J Chem Inf Model 62, 5938–5951 (2022).

6. Li, X. et al. SDNN-PPI: self-attention with deep neural network effect on protein-protein interaction prediction. BMC Genomics 23, 474 (2022).

7. Sledzieski, S., Singh, R., Cowen, L. & Berger, B. D-SCRIPT translates genome to phenome with sequence-based, structure-aware, genome-scale predictions of protein-protein interactions. Cell Syst 12, 969–982 e966 (2021).

8. Yao, Y., Du, X., Diao, Y. & Zhu, H. An integration of deep learning with feature embedding for protein-protein interaction prediction. PeerJ 7, e7126 (2019).

9. Chen, M. et al. Multifaceted protein-protein interaction prediction based on Siamese residual RCNN. Bioinformatics 35, i305–i314 (2019).

10. Liu, D., et al. PLM-interact: extending protein language models to predict protein-protein interactions. bioRxiv (2024).

11. Zhang, N. et al. MutaBind2: Predicting the Impacts of Single and Multiple Mutations on Protein-Protein Interactions. iScience 23, 100939 (2020).

12. Dehouck, Y., Kwasigroch, J.M., Rooman, M. & Gilis, D. BeAtMuSiC: Prediction of changes in protein-protein binding affinity on mutations. Nucleic Acids Res 41, W333–339 (2013).

13. Liu, X., Luo, Y., Li, P., Song, S. & Peng, J. Deep geometric representations for modeling effects of mutations on protein-protein binding affinity. PLoS Comput Biol 17, e1009284 (2021).

14. Wang, M., Cang, Z. & Wei, G.W. A topology-based network tree for the prediction of protein-protein binding affinity changes following mutation. Nat Mach Intell 2, 116–123 (2020).

15. Xiong, D. et al. A structurally informed human protein-protein interactome reveals proteome-wide perturbations caused by disease mutations. Nat Biotechnol (2024).

16. Brandes, N., Goldman, G., Wang, C.H., Ye, C.J. & Ntranos, V. Genome-wide prediction of disease variant effects with a deep protein language model. Nat Genet 55, 1512–1522 (2023).

17. Lin, Z. et al. Evolutionary-scale prediction of atomic-level protein structure with a language model. Science 379, 1123–1130 (2023).

18. Pokharel, S., Pratyush, P., Heinzinger, M., Newman, R.H. & Kc, D.B. Improving protein succinylation sites prediction using embeddings from protein language model. Sci Rep 12, 16933 (2022).

19. Hayes, T. et al. Simulating 500 million years of evolution with a language model. Science 387, 850–858 (2025).

20. Cheng, J. et al. Accurate proteome-wide missense variant effect prediction with AlphaMissense. Science 381, eadg7492 (2023).

21. Liang, S. et al. A sequential recommendation method using contrastive learning and Wasserstein self-attention mechanism. PeerJ Comput Sci 11, e2749 (2025).

22. Wang, Y., Li, Y., Liu, L., Hong, P. & Xu, H. Advancing Drug Discovery with Enhanced Chemical Understanding via Asymmetric Contrastive Multimodal Learning. J Chem Inf Model (2025).

23. Song, Y., Wang, Y., He, H. & Gao, X. Recognizing Natural Images From EEG With Language-Guided Contrastive Learning. IEEE Trans Neural Netw Learn Syst PP (2025).

24. Cao, Z. et al. Decentralized learning for medical image classification with prototypical contrastive network. Med Phys 52, 4188–4204 (2025).

25. Li, W. et al. scPROTEIN: a versatile deep graph contrastive learning framework for single-cell proteomics embedding. Nat Methods 21, 623–634 (2024).

26. Lin, W., Miller, D., Gu, Z. & Orengo, C. GOBeacon: An ensemble model for protein function prediction enhanced by contrastive learning. Protein Sci 34, e70182 (2025).

27. Wang, D. et al. S-PLM: Structure-Aware Protein Language Model via Contrastive Learning Between Sequence and Structure. Adv Sci (Weinh) 12, e2404212 (2025).

28. Du, X. et al. DeepPPI: Boosting Prediction of Protein-Protein Interactions with Deep Neural Networks. J Chem Inf Model 57, 1499–1510 (2017).

29. Chen, C., Zhang, Q., Ma, Q. & Yu, B. LightGBM-PPI: Predicting protein-protein interactions through LightGBM with multi-information fusion. Chemometrics and Intelligent Laboratory Systems 191 (2019).

30. Zhang, L., Yu, G., Xia, D. & Wang, J. Protein-protein interactions prediction based on ensemble deep neural networks. Neurocomputing 324, 10–19 (2019).

31. Fragoza, R. et al. Extensive disruption of protein interactions by genetic variants across the allele frequency spectrum in human populations. Nat Commun 10, 4141 (2019).

32. Miosge, L.A. et al. Comparison of predicted and actual consequences of missense mutations. Proc Natl Acad Sci U S A 112, E5189–5198 (2015).

33. Bryant, P., Pozzati, G. & Elofsson, A. Improved prediction of protein-protein interactions using AlphaFold2. Nat Commun 13, 1265 (2022).

34. Jumper, J. et al. Highly accurate protein structure prediction with AlphaFold. Nature 596, 583–589 (2021).

35. Evans, R., et al. Protein complex prediction with AlphaFold-Multimer. bioRxiv (2021).

36. Clarke, N.F. et al. Mutations in TPM3 are a common cause of congenital fiber type disproportion. Ann Neurol 63, 329–337 (2008).

37. Laing, N.G. et al. A mutation in the alpha tropomyosin gene TPM3 associated with autosomal dominant nemaline myopathy. Nat Genet 9, 75–79 (1995).

38. McArdle, A. et al. HSF expression in skeletal muscle during myogenesis: implications for failed regeneration in old mice. Exp Gerontol 41, 497–500 (2006).

39. Mollet, J. et al. CABC1 gene mutations cause ubiquinone deficiency with cerebellar ataxia and seizures. Am J Hum Genet 82, 623–630 (2008).

40. Lee, M.S. & Olson, M.A. Calculation of absolute protein-ligand binding affinity using path and endpoint approaches. Biophys J 90, 864–877 (2006).

41. Schymkowitz, J.W. et al. Prediction of water and metal binding sites and their affinities by using the Fold-X force field. Proc Natl Acad Sci U S A 102, 10147–10152 (2005).

42. Bernardi, R.C., Melo, M.C.R. & Schulten, K. Enhanced sampling techniques in molecular dynamics simulations of biological systems. Biochim Biophys Acta 1850, 872–877 (2015).

43. Burley, S.K. et al. RCSB Protein Data Bank (RCSB.org): delivery of experimentally-determined PDB structures alongside one million computed structure models of proteins from artificial intelligence/machine learning. Nucleic Acids Res 51, D488–D508 (2023).

44. Fowler, D.M. & Fields, S. Deep mutational scanning: a new style of protein science. Nat Methods 11, 801–807 (2014).

45. Baek, M. et al. Accurate prediction of protein structures and interactions using a three-track neural network. Science 373, 871–876 (2021).

46. Yang, X. et al. A public genome-scale lentiviral expression library of human ORFs. Nat Methods 8, 659–661 (2011).

47. Huang, B. et al. Protein Structure Prediction: Challenges, Advances, and the Shift of Research Paradigms. Genomics Proteomics Bioinformatics 21, 913–925 (2023).

48. Goldman, M.J. et al. Visualizing and interpreting cancer genomics data via the Xena platform. Nat Biotechnol 38, 675–678 (2020).

